# High-performance analysis of biomolecular containers to measure small-molecule transport, transbilayer lipid diffusion, and protein cavities

**DOI:** 10.1101/701573

**Authors:** Alexander J. Bryer, Jodi A. Hadden, John E. Stone, Juan R. Perilla

## Abstract

Compartmentalization is a central theme in biology. Cells are composed of numerous membrane-enclosed structures, evolved to facilitate specific biochemical processes; viruses act as containers of genetic material, optimized to drive infection. Molecular dynamics simulations provide a mechanism to study biomolecular containers and the influence they exert on their environments; however, trajectory analysis software generally lacks knowledge of container interior versus exterior. Further, many relevant container analyses involve large-scale particle tracking endeavors, which may become computationally prohibitive with increasing system size. Here, a novel method based on 3-D ray casting is presented, which rapidly classifies the space surrounding biomolecular containers of arbitrary shape, enabling fast determination of the identities and counts of particles (e.g., solvent molecules) found inside and outside. The method is broadly applicable to the study of containers and enables high-performance characterization of properties such as solvent density, small-molecule transport, transbilayer lipid diffusion, and topology of protein cavities. The method is implemented in VMD, a widely used simulation analysis tool that supports personal computers, clouds, and parallel supercomputers, including ORNL’s Summit and Titan and NCSA’s Blue Waters, where the method can be employed to efficiently analyze trajectories encompassing millions of particles. The ability to rapidly characterize the spatial relationships of particles relative to a biomolecular container over many trajectory frames, irrespective of large particle counts, enables analysis of containers on a scale that was previously unfeasible, at a level of accuracy that was previously unattainable.

**Author summary:** The cell is the basic unit of life. Within the container of the cell, the many chemical reactions and biological processes essential to life are carried out simultaneously. Human and other eukaryotic cells include a variety of sub-containers, namely organelles, that provide separation between reactions and processes, and engender the chemical environments conducive to them. In order to understand how the cell works, researchers must study the functions of these containers. Molecular dynamics simulations can reveal important information about how biomolecular containers behave and control their enclosed environments, but the latter can be particularly challenging and expensive to measure. The challenge arises because simulation analysis software lacks awareness of the concepts of container “inside” and “outside.” The expense arises because tracking the many solvent molecules that make up a container’s environment requires significant computing power. We have developed a method that allows the simulation analysis software VMD to automatically detect the interior versus exterior of a container and quickly identify the solvent molecules found in each location. This versatile new feature enables researchers to characterize essential container properties using a relatively inexpensive calculation. Further, the method performs efficiently on supercomputers, allowing researchers to study massive container systems that include millions of particles.

## Introduction

Living cells are complex containers, within which numerous reactions and biochemical processes must be carried out simultaneously to support life. Eukaryotes partition their intracellular space with sub-containers, membrane-enclosed structures including organelles and vesicles, allowing them to organize and regulate the various aspects of their metabolism. Viruses are also containers, packaging their genomes and accessory enzymes in protein shells called capsids, which may be further wrapped in lipid envelopes. Proteins themselves may include cavities embedded in their tertiary or quarternary structures, which can act as containers for ligands or substrates that recognize and bind them.

Importantly, all biomolecular containers are highly specialized to facilitate the specific interaction, reaction, or process they enclose, and exhibit key biophysical properties that enable them to carry out their functions. Protein cavities that serve as binding sites engender ligand affinity by offering suitable pocket size and shape complementarity. Vesicles and capsids often play essential roles in maintaining the biochemical environments necessary for their compartmentalized processes, regulating the concentration, pH, or pressure of their interior space by controlling the passage of small molecules, ions, protons, or solvent. Some membrane-bound structures must establish or preserve asymmetry—differing lipid compositions across the inner and outer leaflets of the bilayer—to drive cellular functions, requiring transbilayer diffusion and protein-catalyzed translocation of lipids between leaflets.

Molecular dynamics (MD) simulations are now routinely applied to study large biological systems, including containers such as virus capsids and viral envelopes, at both coarse-grained and all-atom levels of detail [1–3]. Because MD simulations can be used to characterize the influence of containers on their surrounding biochemical environments at the resolution of individual particles—a level of detail inaccessible to experimental methods—the technique provides a powerful approach to investigate the properties of containers that underlie their functions. For example, by tracking the exchange of solvent and ions across the surface of virus capsids, researchers have been able to determine their small-molecule transport properties, revealing insights into the capsids’ regulatory roles in reverse transcription and cellular signaling [4, 5]. Due to the size of virus capsids and most other containers of biological interest, however, simulated systems can encompass millions of particles, with those describing the solvent environment far outnumbering solute.

In the present manuscript, a novel method is presented that enables rapid analysis of biomolecular containers, irrespective of large system particle counts. The method, implemented in VMD [6] version 1.9.4 and referred to herein by its VMD command *measure volinterior*, automatically distinguishes container interior versus exterior, regardless of container shape, and detects the spatial relationships of particles relative to the container in a manner that is efficient and highly-scalable. Instead of dealing with particles individually, *measure volinterior* uses 3-D ray casting to classify the space surrounding the container’s molecular surface. VMD’s atom selection feature then permits rapid collection of the identities of particles that are located inside or outside of the container. This information can be used to determine the number and density of particles within or around a container, characterize small-molecule transport by computing the exchange rates of particles flowing inward and outward across the container’s surface, track transbilayer diffusion in lipid-based containers, and even identify and measure protein cavities that may represent small-molecule binding pockets or druggable sites.

In the following sections, details of *measure volinterior* theory and implementation are provided, along with guidelines for selecting appropriate parameters to define the molecular surfaces of containers required for the method. Code examples are also included to demonstrate the application of *measure volinterior* to perform the container analyses listed above. Finally, the protocol for analyzing an MD simulation trajectory of a multimillion-atom biomolecular container is described, taking advantage of supercomputers such as ORNL’s Titan and Summit and NCSA’s Blue Waters, which include access to high-performance Lustre file systems.

## Methods

### Theory and implementation of *measure volinterior*

A tractable approach for analyzing MD simulations of biomolecular containers requires automated execution of the following in an efficient and highly-scalable manner to enable rapid processing of millions of particles over thousands of trajectory frames: i. recognition of the container, ii. detection of container interior versus exterior, regardless of container shape or orientation, and iii. classification of non-container particles according to their location inside or outside of the container. The *measure volinterior* approach addresses these challenges by using a 3-D ray casting method combined with a spatially discretized or *voxelized* system and the space around it. Ray casting has previously been proposed as a general and effcient strategy for characterizing properties of molecules that arise from their surface geometry [7]. Importantly, *measure volinterior* can be applied to study containers of arbitrary shape, and the results it produces are spatially invariant. The algorithm is implemented in C++ in VMD [6] version 1.9.4, where ray-level parallelism maximizes computational performance.

The VMD command for calling *measure volinterior* on a container described by atom selection *sel* in molecule *mol*, is given in Code Snippet 1. A given container is defined by its molecular surface, calculated using VMD’s QuickSurf algorithm. The user must provide parameters for *Radius Scale, Isovalue*, and *Grid Spacing*. For accurate results, the parameters must yield a nonporous container, characterized by a continuous molecular surface without holes; otherwise the user must enable fuzzy-boundary detection. See the following sections for guidelines for selecting QuickSurf parameters for various types of biomolecular containers, as well as details on dealing with fuzzy container boundaries. The user must also indicate the number of rays, *n*. Upon calling *measure volinterior* on a container selection, a 3-D grid is generated with the same dimensions as the system bounding box for the given trajectory frame. For each voxel in the grid, *n* rays are cast outward. In the standard *measure volinterior* implementation, the container interior versus exterior is determined based on whether the rays strike the system bounding box or are blocked by the container wall (Fig. 1A).

**Code Snippet 1.**
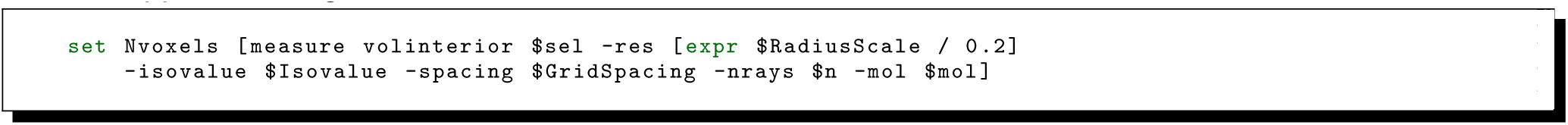
Calling *measure volinterior* on a container selection.

**Fig 1.**
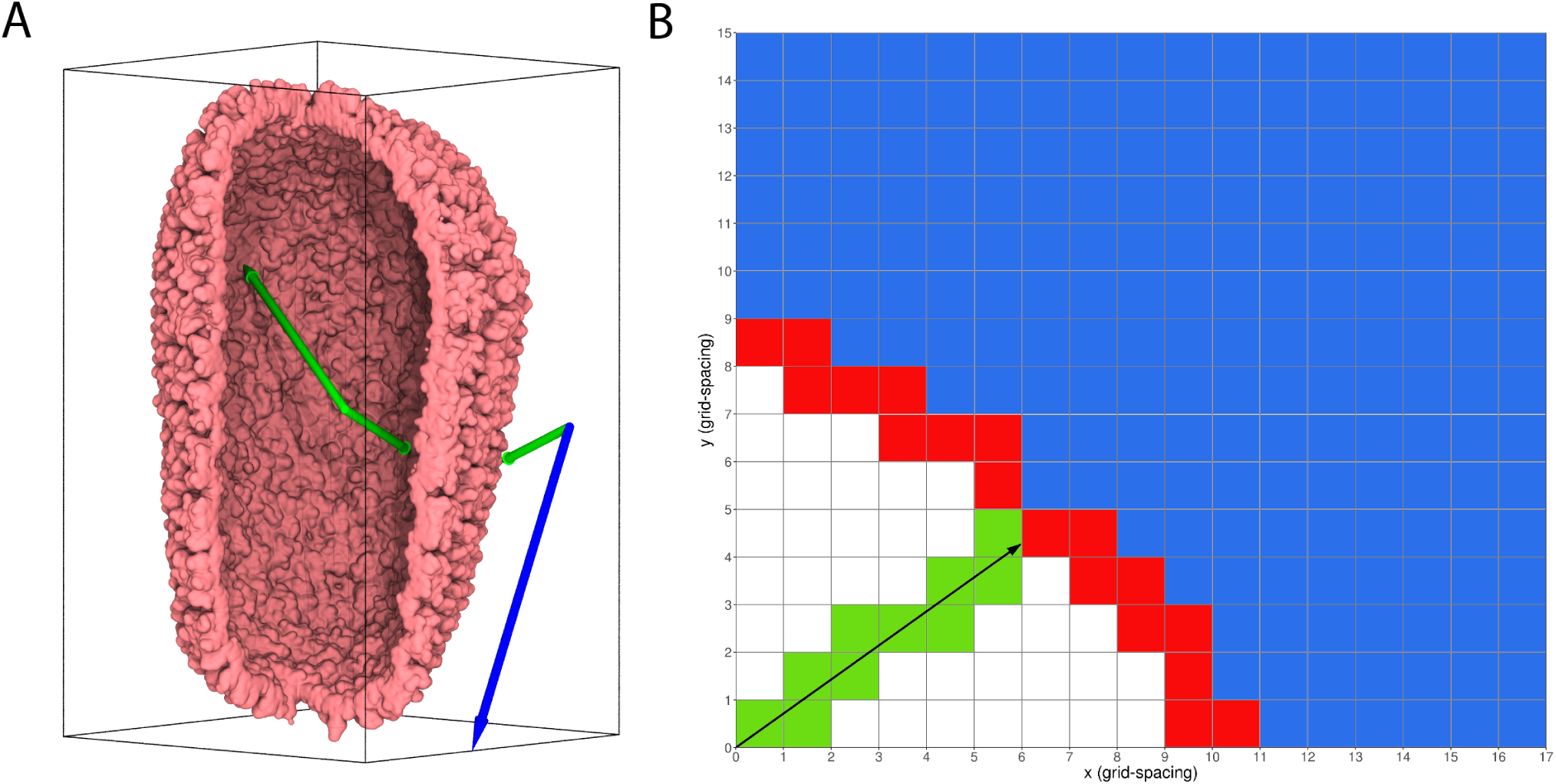
Ray casting and voxel traversal. (A) Schematic representation of *measure volinterior* applied to a cross-section of the HIV-1 capsid. Here, *n*=2 rays are cast from each voxel in a grid set to the size of the system bounding box. The rays are randomly and uniformly distributed, according to the spherical distribution given in Eq. 1. For a given voxel shown on the capsid’s exterior, one of the rays (blue) traverses the grid of voxels unimpeded over the entire interval up to the wall of the bounding box, indicating that the voxel is located outside the capsid container. For a given voxel shown on the capsid’s interior, both of the rays (green) fail to strike the wall of the bounding box because they intersect (are blocked by) the capsid surface, indicating that the voxel is located inside the capsid container. (B) A 2-D version of the DDA algorithm. The ray, cast from the grid origin, traverses the grid visiting a sequence of cells (equivalent to voxels in 3-D) on its path to the system’s bounding box. In this example case, the ray strikes voxels occupied by the container (red), determining that the ray originated inside the container (green) instead of outside (blue).

Each ray is cast through a 3-D grid defined by the system bounding box, using the Digital Differential Analysis (DDA) algorithm [8]. The DDA voxel traversal algorithm incrementally computes x, y, and z voxel indices, visiting all voxels pierced by the ray between its origin and a wall or obstacle. Using the DDA algorithm for voxel traversal, the rays are rasterized passing through the aforementioned voxelized space, sampling the voxel contents to check for container intersections, thereby allowing the starting voxel to be classified as interior or exterior. A 2-D version of the algorithm is illustrated in Fig. 1B, depicting the process of voxel traversal by a ray and the resulting classification of voxel space.

In DDA, rays are cast outward from the origin of each voxel with directions following a random, uniform spherical distribution given by

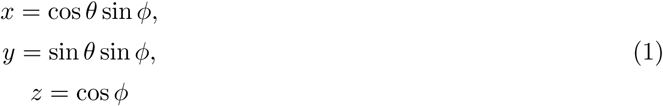

where

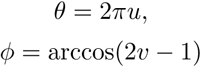

and *u* and *v* are uniformly distributed on [0, 1], that is

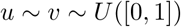

The direction (*x, y, z*) is then used to compute a step along the tracing ray, such to match the width of one voxel in each dimension, as follows

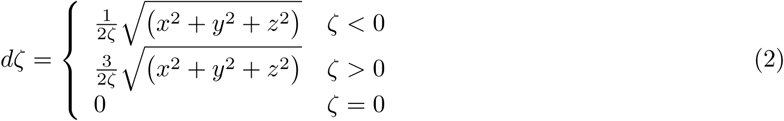

where

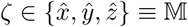

and where 𝕄 is the set of all unitary vectors.

With the differentials defined by Eq. (2), DDA determines digital steps of size *m*_*ζ*_ that propagate the ray in voxel space, one voxel index incremement at a time. To accomplish the ray propagation, the algorithm first finds the minimum valued *ζ* of the current direction vectors. The minimum, or lagging dimension is given a step *m*_*ζ*_ of ±1 and the other two dimensions are given a step of 0. Therefore, the ray propagates according to

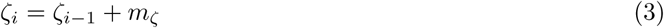

where *i* = 1 *… N*. *N* is determined conditionally based on whether the ray intersects voxels belonging to the molecular surface or strikes the bounding box.

If any one of the rays cast from a voxel strikes the bounding box, then that voxel is considered to be located on the container’s exterior and marked with a grid value of 5.0 (Fig. 1A, blue arrow). Conversely, if none of the rays cast from a voxel strike the bounding box, because they are all blocked by the inner walls of the container, then that voxel is considered to be located on the container’s interior and marked with a grid value of 0.0 (Fig. 1A, green arrows). Voxels that coincide spatially with the container surface and represent the boundary between interior and exterior are considered container voxels, marked with a grid value of −5.0,

The output of *measure volinterior* is a list containing the total voxel count and the number of voxels identified as inside, outside, or comprising the container, as well as a 3-D grid map (e.g., vol0). The voxels of the map are assigned with values indicating their interior (vol0 < 1 and vol0 > −1), exterior (vol0 > 0), or boundary (vol0 < 0) classification for the processed trajectory frame. The numbers of voxels identified in each region can be accessed via their list indices, as shown in Code Snippet 2. Further, by invoking the corresponding grid values from the 3-D map, VMD’s atom selection feature allows for fast region classification of system particles according to their location. Particles that happen to fall outside of the system bounding box, whose associated space is not detected by *measure volinterior*, likely also fall on the container exterior and their map values can be selected as those that are not a number (vol0 > 1 and vol0 < −1). Importantly, the relational operators used for atom selections should be written as intervals instead of equalities (e.g., vol0 < 1 and vol0 > −1 instead of vol0 == 0) to ensure platform independent behavior. Code Snippet 3 shows how to define macros for classified regions of the output map, for use in atom selections.

**Code Snippet 2.**
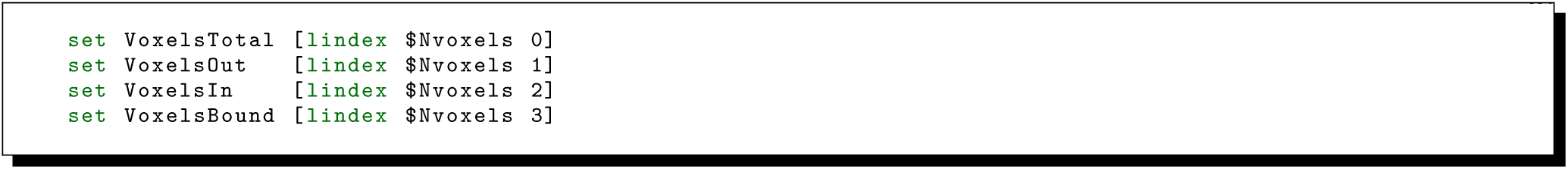
Accessing indexed voxel counts returned by *measure volinterior*.

**Code Snippet 3.**
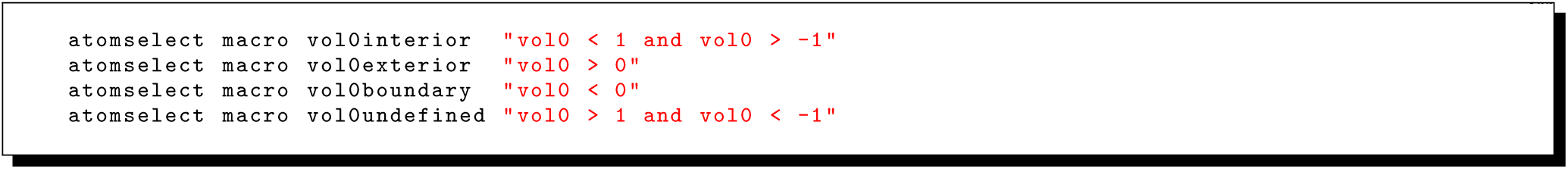
Assigning atom selection macros for use with *measure volinterior*.

VMD 1.9.4, which includes full *measure volinterior* functionality, is compiled on ORNL’s Titan and Summit and NCSA’s Blue Waters supercomputers, and is readily available to any researcher with access to these resources for post-processing analysis of large MD simulation trajectories. For smaller systems, the method also performs well on desktop workstations. See the **Results and Discussion** section for examples demonstrating the usage of *measure volinterior* to analyze biomolecular containers for properties such as enclosed solvent density, small-molecule transport, and transbilayer lipid diffusion, as well as to detect and measure cavities within or between constituent proteins.

### Selecting parameters to define container surfaces

When using *measure volinterior*, containers are defined by their molecular surfaces, as calculated by VMD’s QuickSurf algorithm. QuickSurf generates a molecular surface extracted from a volumetric Gaussian density map, computed by summing the Gaussian densities from the particles in a region surrounding each grid point [9,10]. User-defined parameters allow manipulation of the molecular surface representation to alter spatial fidelity and level of detail. For the standard implementation of *measure volinterior*, accurate characterization of a container and its biophysical properties depends on selecting QuickSurf parameters that yield a continuous molecular surface with no holes. Otherwise, fuzzy-boundary detection must be enabled (see below). In the standard case, surface defects and indentations do not impact the performance of *measure volinterior*, as long as they do not penetrate the surface completely. Overall, the container surface must be nonporous, such that rays cast from inside the container do not inadvertently escape to the container exterior, resulting in incorrect classification of the surrounding space. This means that containers with natural pores or channels, such as virus capsids, require QuickSurf parameters designed to mask such openings without sacrificing other important features of the container surface.

Three QuickSurf parameters are required by *measure volinterior* to define molecular surfaces, namely *Radius Scale, Isovalue*, and *Grid Spacing. Radius Scale* is a radius scaling factor applied to all particles prior to computing their contributions to the volumetric density map; its value relates to overall spatial resolution of the final surface. A larger value of *Radius Scale* will produce a bulkier molecular surface, which may be needed to mask pores and prevent the escape of rays to the container exterior. *Isovalue* is the isovalue used when extracting the molecular surface from the density map and represents the scalar density with which the surface is generated. *Grid Spacing* is the distance between grid points, which defines the size and number of voxels. Smaller values of *Grid Spacing* will resolve finer details in the molecular surface; a value of 1 Å is sufficient to represent the surface at atomic detail. For larger container systems, it may be desirable to use larger values of *Grid Spacing*, sacrificing surface resolution for computational efficiency. Since a lower resolution grid contains fewer voxels, and *n* rays are cast per voxel, larger values of *Grid Spacing* result in significantly fewer rays cast per processed trajectory frame. Further, a lower resolution grid entails a smaller memory footprint and may prevent memory overflow during analysis of a large container. Decreasing the number of rays can also improve computational efficiency. While *n*=64 rays will likely be sufficient to characterize a highly detailed molecular surface, *n*=32 rays may represent a reasonable compromise for large container systems, especially if their surfaces are relatively featureless.

Selection of suitable inputs for *measure volinterior* is an empirical process. VMD’s graphical interface should be used to experiment with and identify parameters for QuickSurf that capture the essential topology of the biomolecular container of interest. For use with the standard implementation of *measure volinterior*, the parameters should yield a molecular surface that is continuous. The output of *measure volinterior* for a given trajectory frame should be visualized to assess the performance of a candidate parameter set prior to production analysis. Fig. 2 shows cross-sections of several container systems, along with volume slices from 3-D grid maps produced by calling *measure volinterior* on them. The parameters used to generate the maps, provided in Table 1, capture significant detail in the containers’ surface features, while successfully masking pores and channels to prevent the escape of rays. The resulting classification of space around the containers is highly accurate, with the delineations of interior versus exterior conforming closely to the molecular structure.

**Table 1.**
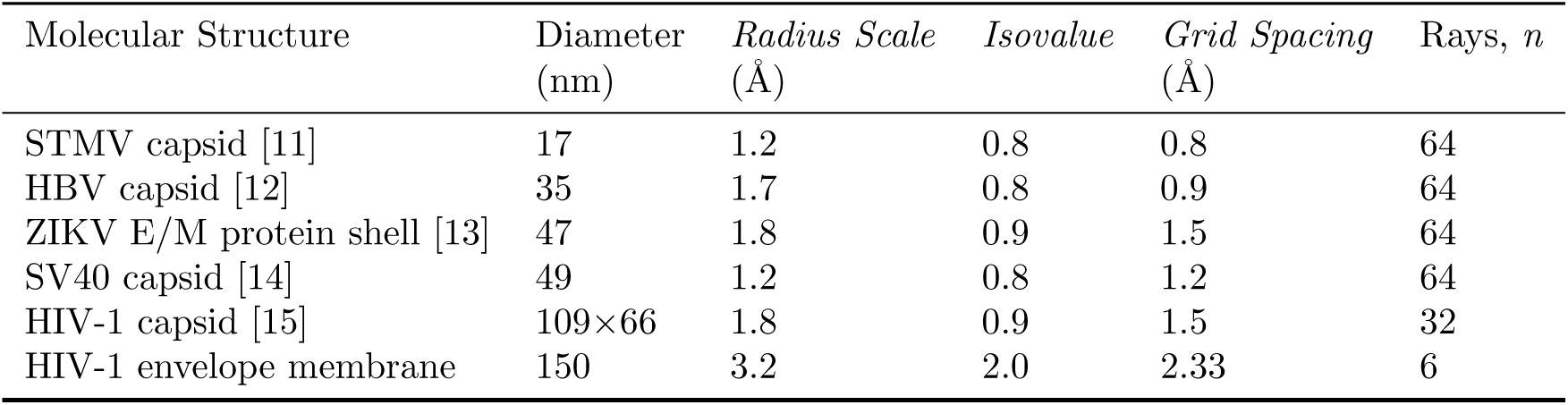
*measure volinterior* parameters used to generate the 3-D grid maps shown in Fig. 2.

**Fig 2.**
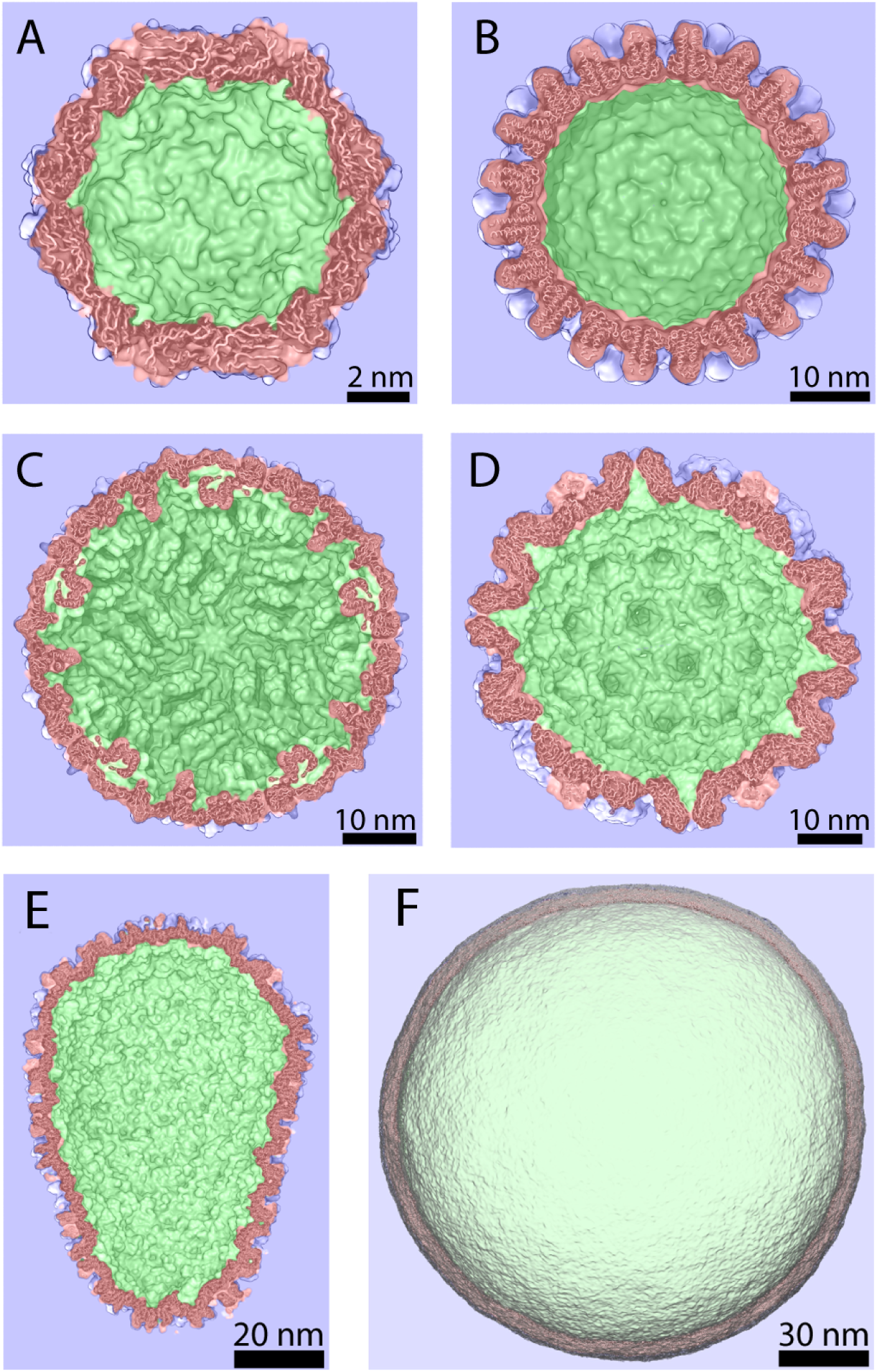
Visualizing the output of *measure volinterior*. Six biomolecular container structures are shown in cross-section, along with a volume slice from the 3-D grid map produced by *measure volinterior* (e.g., vol0). Using the parameters listed in Table 1, space classification as interior (vol0interior, green), exterior (vol0exterior, blue), or boundary (vol0boundary, red) closely matches the molecular surface, capturing notable topological detail. See Code Snippet 3 to see how the macros specifying these regions of the map, e.g., vol0interior and vol0exterior, are defined. (A) satellite tobacco mosaic virus (STMV) capsid [11], (B) hepatitis B virus (HBV) capsid [12], (C) zika virus (ZIKV) envelope/membrane (E/M) protein shell [13], (D) simian vacuolating virus 40 (SV40) capsid [14], (E) HIV-1 capsid [15], and (F) HIV-1 envelope membrane.

Fig. S1 illustrates the effects of using parameters that are not well-suited to produce accurate results with *measure volinterior*. Importantly, parameters selected for a given system must perform well for all frames of a simulation trajectory. Because pores and channels may fluctuate in diameter during the course of simulation, parameters selected for use with the standard implementation of *measure volinterior* must produce a molecular surface sufficiently thick to mask openings over all sampled conformations. For example, QuickSurf parameters optimized for the experimental HBV capsid structure (Fig. 2B) allow the escape of rays to the container exterior for some frames during MD simulation at 310 K [5] (Fig. S1A, left); slight adjustments to the parameters produce a closed surface and ensure accuracy over all frames during production trajectory analysis (Fig. S1A, right). On the other hand, parameters that produce an overly bulky surface will capture less topological detail and also lead to more space surrounding the container being classified as part of the container itself. For example, QuickSurf parameters that resolve fine surface definition in the ZIKV E/M protein shell lead to highly accurate classification of the space surrounding the container (Fig. S1B, left); a bulkier surface leads to extra space adjacent to the container being counted as part of the boundary (Fig. S1B, right).

### Dealing with fuzzy container boundaries

To analyze containers with discontinuous molecular surfaces, such as those that include pores or holes, or those that represent open cavities, *measure volinterior* is equipped with an algorithm for fuzzy-boundary detection. In the standard *measure volinterior* implementation, if any ray cast from a voxel strikes the sytem bounding box, that voxel is classified as being outside of the container; the output map contains discrete space assignments of inside, outside, or boundary between (i.e., the container itself). When fuzzy-boundary detection is enabled, *measure volinterior* assigns a probabilty to each voxel based on the number of rays cast from it that fail to strike the bounding box. The probability describes, generally, the extent to which a voxel is occluded by the molecular surface or the degree to which it represents a cavity region. In that case, a probability threshold or interval can be used to define the boundary between container interior and exterior, instead of relying on a closed shell to delineate these regions. Fig. 3 shows cross-sections of two container systems that contain pores or holes, along with volume slices from 3-D grid maps produced by calling *measure volinterior* on them with fuzzy-boundary detection enabled. Despite discontinuous molecular surfaces, the fuzzy-boundary detection method accurately identifies the interior versus exterior regions of the containers.

**Fig 3.**
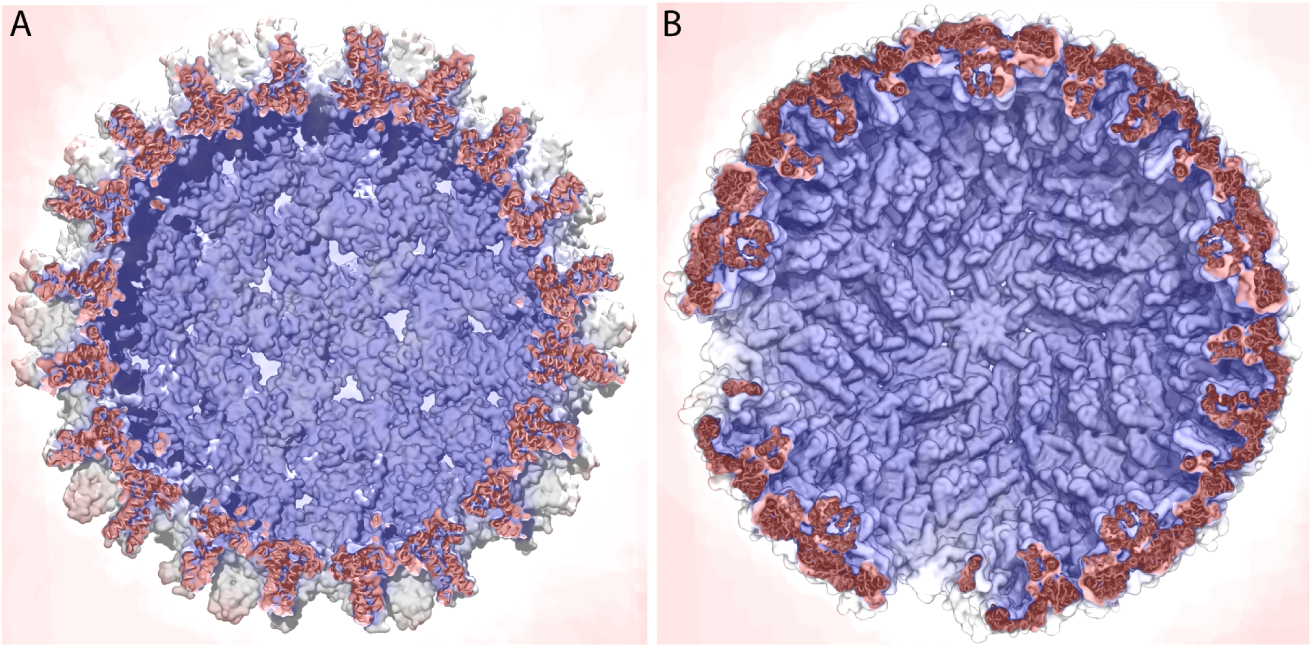
Visualizing the output of *measure volinterior* with fuzzy-boundary detection. Two biomolecular container structures are shown in cross-section, along with a volume slice from the 3-D grid map produced by *measure volinterior* with fuzzy-boundary detection (e.g., vol1). (A) The pores of the HBV capsid [12] must be masked with bulky QuickSurf parameters for accurate analysis with the standard implementation of *measure volinterior*. The fuzzy-boundary detection method accurate identifies the capsid interior versus exterior despite the presence of pores. (B) Even with entire subunits removed from the ZIKV E/M protein shell [13], the fuzzy-boundary detection method is able to delineate its interior versus exterior based on the extent to which these regions are occluded by the molecular surface.

Fuzzy boundary detection is based on the standard implementation of *measure volinterior*, but includes changes to the conditional logic used to assign voxel values. While the standard implementation employs a break condition to halt ray casting for a given voxel after the first of *n* rays strike the system bounding box, (which minimizes computational expense), fuzzy-boundary detection requires that the full set of *n* rays are cast. Subsequently, a 3-D grid *T* stores values between [0, *n*], corresponding to the number of rays from a given voxel that intersect the molecular surface. The probability map *P* is computed according to

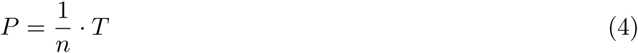

where values in the map range between [0, 1] and *P* = 1 describes a completely occluded voxel.

The VMD command for calling *measure volinterior* with fuzzy-boundary detection is given in Code Snippet 4. The probabilities calculated for each voxel using fuzzy-boundary detection will depend on the total number of rays *n*. By invoking grid values from the output map (e.g., vol1), users can use probability thresholds or intervals in their atom selections to define the boundary between container interior and exterior and classify particles according to their respective locations. Just as suitable QuickSurf parameters must be determined empirically for each container system based on visualization of *measure volinterior* results, the probability thresholds or intervals used with fuzzy-boundary detection must also be experimented with to determine what represents the ideal for each system prior to production analysis. Code Snippet 5 shows examples of how to define macros for regions of the output map, for use in atom selections; the examples pertain to a hypothetical System X, for which the given values were deemed appropriate following visual assessment.

**Code Snippet 4.**
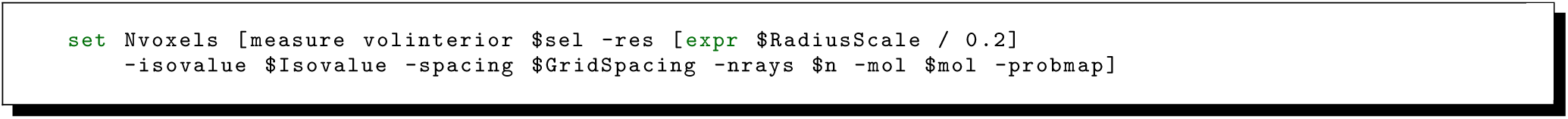
Calling *measure volinterior* with fuzzy-boundary detection.

**Code Snippet 5.**
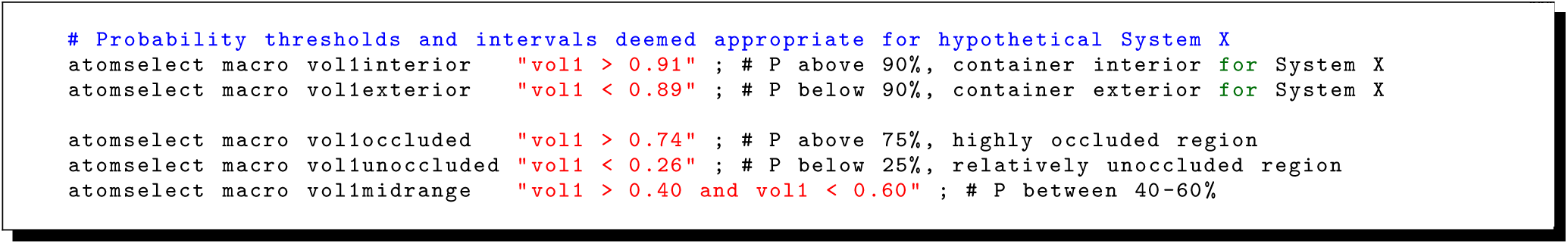
Assigning atom selection macros for use with *measure volinterior* fuzzy-boundary detection.

## Results and Discussion

### Using *measure volinterior* to analyze containers

The fast and accurate detection of a biomolecular container’s interior versus exterior by *measure volinterior* extends the ability of VMD’s atom selection feature to collect the numbers and identities of particles that fall within these regions. The capability to efficiently track particle counts and locations over a series of trajectory frames enables a number of relevant container analyses, such as those presented below, even for multimillion-atom container systems.

### Particle Densities

Biomolecular containers must regulate the density of particles, particularly water molecules, that they enclose in order to maintain correct solvent density. If the container is relatively impermeable, internal density that is too high or too low can lead to rupture or implosion, respectively. Moreover, tracking of enclosed particle densities represents a critical post-processing assessment of any biomolecular container trajectory, necessary to ensure simulation equilibration and validity. Water density can be readily calculated using the relationship [16] given by

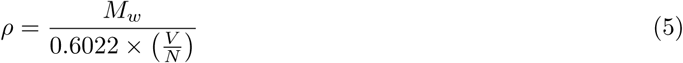

where *M*_*w*_ is the molecular weight of the employed water model (e.g., 18.0154 g *mol*^−1^ for TIP3P), *V* is the volume of the region of interest (i.e., container interior or exterior, in Å^3^, as shown for the HIV-1 capsid in Fig. S2), and *N* is the number of particles located in the given region. A Tcl script demonstrating the measurement of water density within a biomolecular container using *measure volinterior*, is presented in Code Snippet 6.

**Code Snippet 6.**
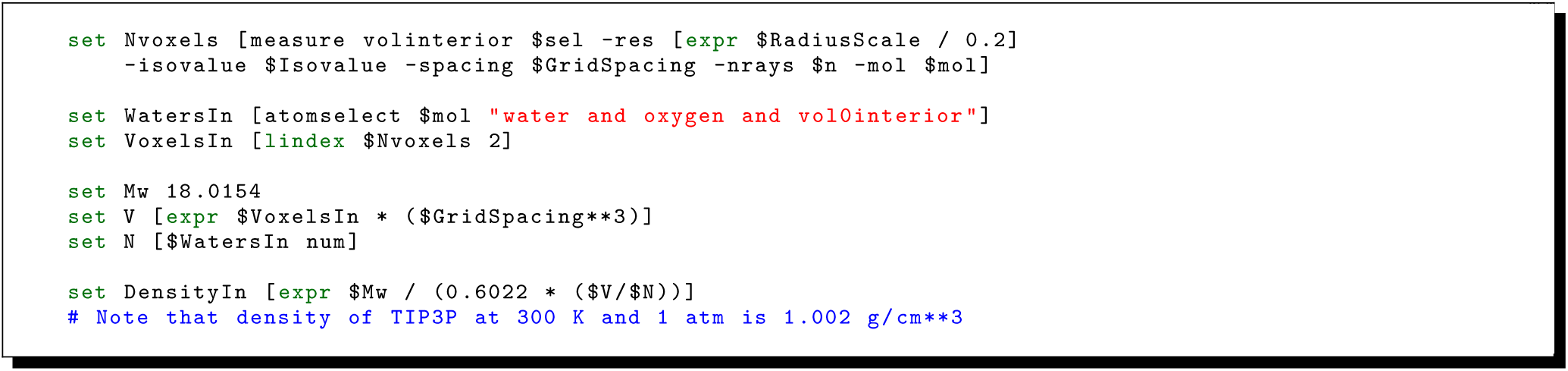
Measuring enclosed water density using *measure volinterior*.

### Small-Molecule Transport

Biomolecular containers often play a role in regulating the biochemical processes they compartmentalize by controlling the passage of small molecules, ions, protons, or solvent [17, 18]. Many such containers act as semi-permeable barriers, allowing the passage of particular species, while selectively filtering others. The transport properties of containers, or the rates at which they exchange molecular species across their surfaces, can be readily calculated by tracking the particles found inside versus outside over time. Notably, such exchange rates can only be obtained using MD simulations, which provide information on the container’s surrounding solvent environment at the resolution of individual particles. For example, rates of solvent exchange have been recently reported for the HIV-1 and HBV capsids based on microsecond MD simulations, revealing that HIV-1 preferentially transports chloride, while HBV preferentially transports sodium [4, 5].

A Tcl script demonstrating an approach to calculate solvent exchange rates using *measure volinterior* is presented in Code Snippet 7 (see Supplement). Exchange is measured based on the identities of solvent species present on one side of the container at a reference frame *t*_0_ and then found on the opposite side of the container at a subsequent frame *t* > *t*_0_ [4, 5, 19]. The cumulative numbers of solvent species transported inward/outward per simulation time segment are exported to plotting software, and the slope of the linear fit to the inward/outward curves gives the respective rates. Examples of such fitting plots are available in the literature [4, 5]. Equivalence between inward and outward exchange rates indicates that the container system has reached equilibrium with its solvent environment. Importantly, the analysis aims to track transport of molecular species from bulk solvent inside the container to bulk solvent outside the container, and vice versa. The accuracy of the calculation may be increased by using a slightly bulkier molecular surface for *measure volinterior*, which expands the boundary region between container interior and exterior and, thus, minimizes detection of recrossing events close to the structure, such as those that might arise when a species tends to localize within a container pore instead of exchanging directly between bulk solvent regions.

### Transbilayer Lipid Diffusion

The transbilayer movement of lipids across membrane-bound containers, commonly referred to as lipid flip-flop, is an essential biological process [20, 21]. This exchange of lipids between the inner/outer leaflets can occur via free diffusion or protein-catalyzed translocation [22]. Flip-flop is an important mechanism for membrane equilibration and enables the regulation of differing lipid compositions across the leaflets of a bilayer, a container property essential to the function of many organelles, including the plasma membrane [23]. Rates of lipid flip-flop can be readily obtained by tracking lipids of a given species found in the inner and outer leaflets over time. Various experimental approaches have been developed to characterize protein-free lipid translocation [24], and transbilayer lipid diffusion can also be observed during MD simulations.

A Tcl script demonstrating the detection of the inner and outer leaflets of a membrane using *measure volinterior*, as well as an approach to track transbilayer diffusion, is presented in Code Snippet 8 (see Supplement). Accurate detection of the inner and outer leaflets requires the parameterization of a molecular surface for *measure volinterior* that corresponds to the structure of the lipid tail domains sandwiched within the membrane bilayer (Fig. 4). Given such a surface, the lipid head groups can be clearly identified as members of the inner and outler leaflets based on their locations inside and outside of the container core, respectively. Exchange between leaflets is measured based on the identities of lipid species present in one leaflet of the bilayer at a reference frame *t*_0_ and then found in the opposite leaflet at a subsequent frame *t* > *t*_0_. Rates of flip-flop observed on the simulation timescale can be determined based on plotting the cumulative numbers of lipid species transported to the inner/outer leaflet per simulation time segment and taking the slope of the linear fit to the respective curves, similar to the analysis of solvent exchange rates discussed above.

**Fig 4.**
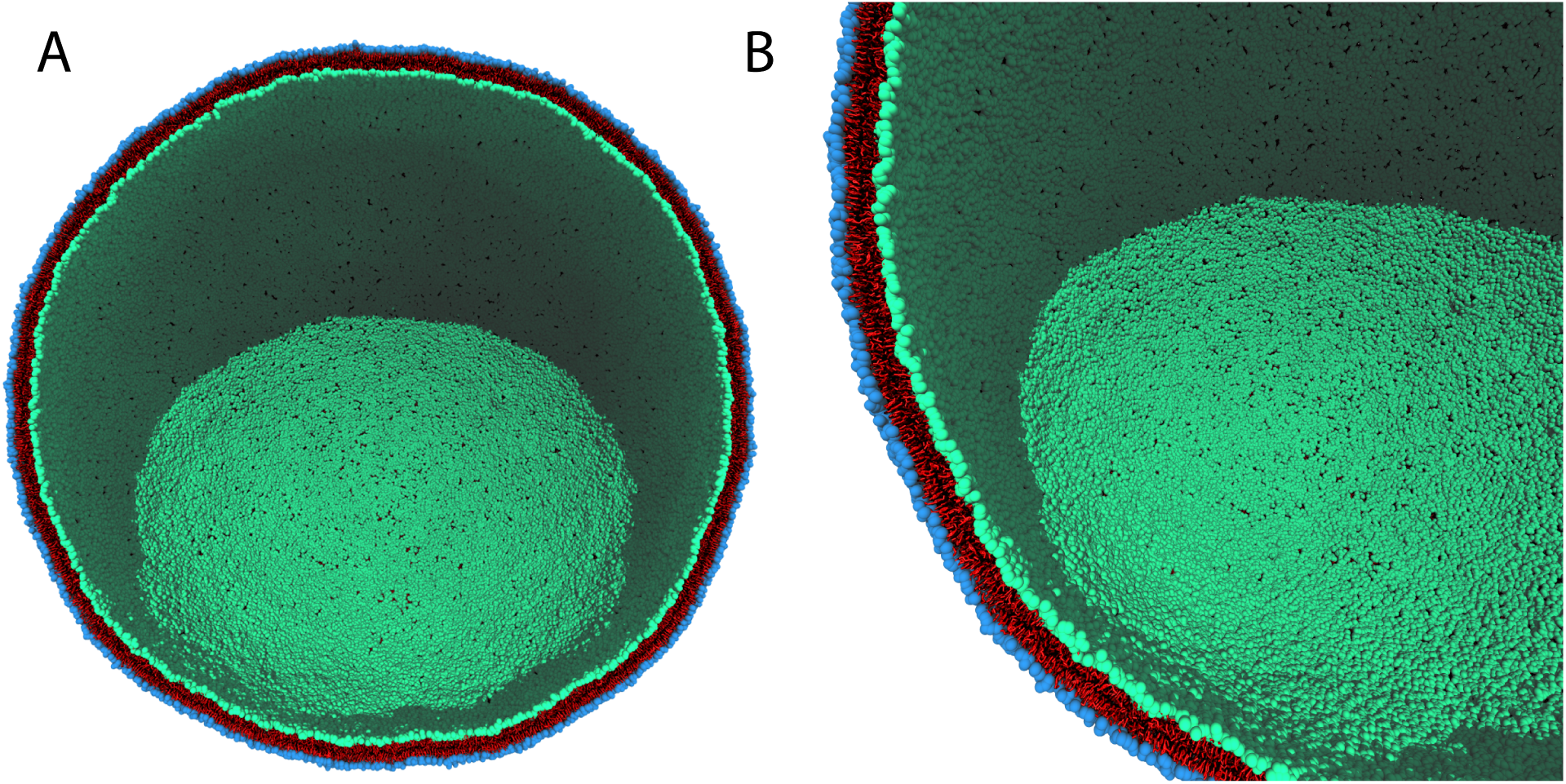
Detection of inner and outer leaflets of a lipid bilayer. A coarse-grained model of the HIV-1 envelope membrane is shown in cross-section in (A) full and (B) quarter views. The parameters listed in Table 1 produce a molecular surface that corresponds to the structure of the lipid tail domains, and *measure volinterior* classifies the associated space as boundary region, colored red. Lipids located on the container interior, identified as members of the inner leaflet, have their head groups colored green; lipids located on the container exterior, identified as members of the outer leaflet, have their head groups colored blue. Leaflet classification is highly accurate, down to level of individual particles.

### Protein Cavities

Cavities embedded within protein structures are often key structural features that contribute to their biomolecular function. These regions my serve as binding sites for native ligands, form the openings of transport channels, or represent potential druggable sites. The identification and characterization of protein cavities is important for the study of fundamental biophysical properties like protein stability, elucidation of the molecular mechanisms underlying cavity-related protein function, and even the design of cavity-targeting therapeutics [25–28]. Numerous computational approaches have been developed to address needs of the latter [29], most aiming to perform some combination of cavity detection and visualization, analysis of cavity topology, volume, flexibility, and/or accessibility, prediction of cavity druggability scores, and/or selection of cavity conformers to generate ensembles for use in high-throughput computational screening of drug compounds. Popular software tools for the examination of cavities include POVME [30–32], fpocket [33, 34], MDpocket [35], TRAPP [36, 37], Epock [38], VolSite [39], and trj cavity [40].

The *measure volinterior* method provides a novel approach for detecting and measuring the spatial properties of protein cavities. Given that cavities are conceptualized essentially as containers, having interior and exterior regions, *measure volinterior* lends itself readily to the task. When protein cavities are buried and represent closed containers, analysis with the standard *measure volinterior* implementation is sufficient to enable their characterization. For example, Fig. 5A shows the results of applying *measure volinterior* to four structural variants of *Staphylococcal* nuclease (wild type, I92A, V66A and L125A); the revealed cavity regions correspond closely to those previously identified for the system. Importantly, the sensitivity of *measure volinterior* to molecular surface detail accurately captures variations in cavity size, shape, and intramolecular location arising from mutations, a biological phenomenon of known significance in drug development [41–43].

**Fig 5.**
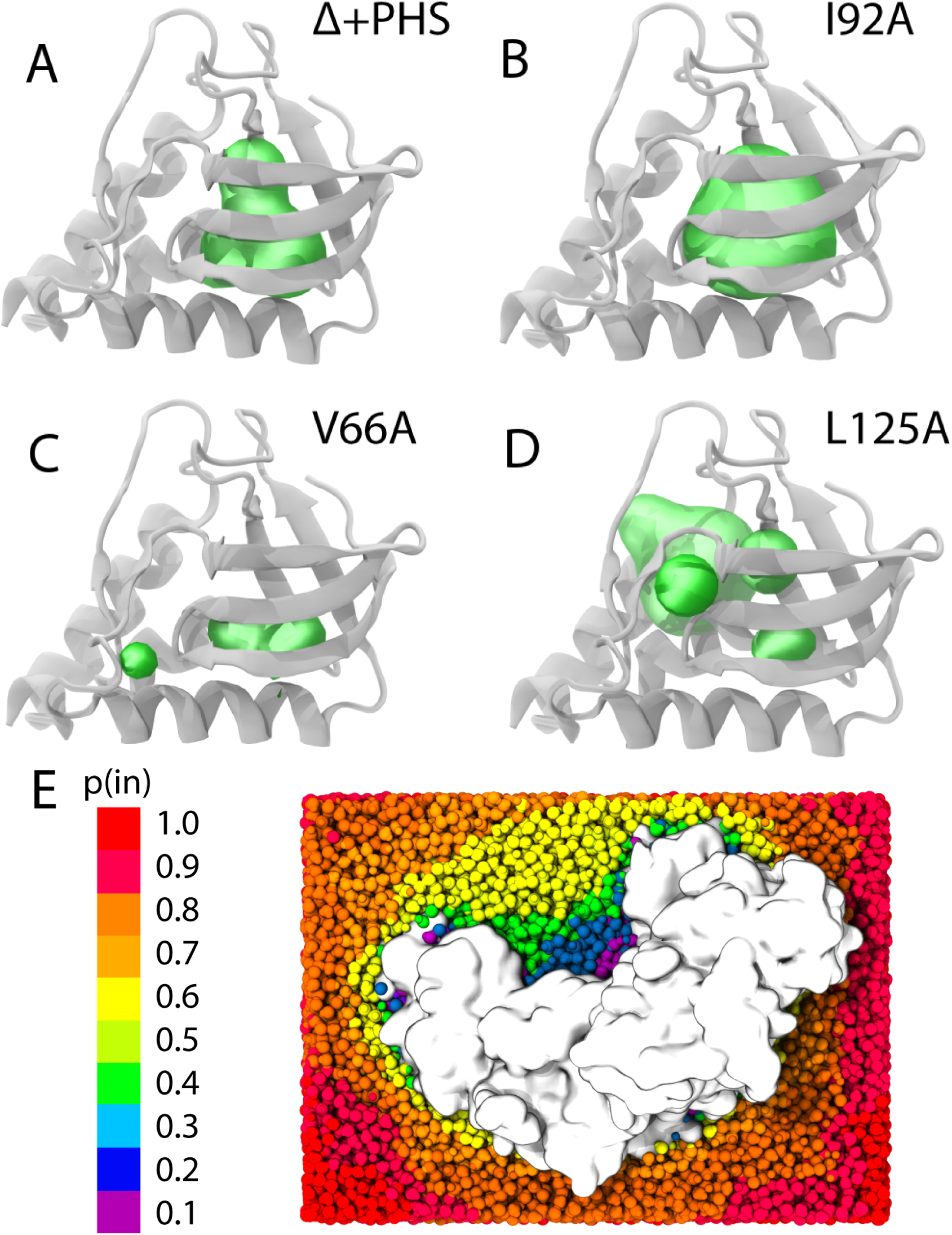
Detection of protein cavities. Four structural mutants of staphylococcal nuclease [44] are shown, with protein cavities identified using the standard implementation of *measure volinterior* highlighted as green surfaces. (A) Δ+PHS wild type, (B) I92A, (C) V66A, and (D) L125A. Cavities are detected as the interior of contained spaces buried within the proteins. Point mutations in the system lead to differences in cavity shape, volume, and intramolecular location. Parameters of *RadiusScale*=0.9, *Isovalue*=0.5, *Grid Spacing* = 1.3, and *n*=12 were used for *measure volinterior*. (E) The N-terminal domain of the glutamate receptor GluA1 [**?**] is shown, immersed in solvent. Solvent molecules are colored according to the output of *measure volinterior* with fuzzy-boundary detection enabled. Probability intervals correspond to the extent to which regions are occluded by the molecular surface and can be used to define the boundary of the cavity.

To enable visualization of the internal volumes of cavities identified by *measure volinterior*, such as shown in Fig. 5A, the *Solvate* plugin in VMD can be used to generate a water box that overlays with the protein structure; by loading the 3-D grid map output by *measure volinterior* into the water box, atom selections applied to water molecules that reference interior space (i.e., vol0interior) appear as 3-D, space-filling representations within the cavity regions. The volume of a protein cavity can be calculated in the same way that the internal volume of a container is calculated in Code Snippet 6.

When protein cavities are solvent exposed and represent open containers, *measure volinterior* ‘s algorithm for fuzzy-boundary detection can be enabled. This approach allows the region that should be considered internal to the cavity to be defined based on the extent to which it is occluded by the molecular surface. In this way, fuzzy-boundary detection provides a novel strategy for handling the long-standing boundary definition problem for open binding pockets [45]. For example, Fig. 5B shows the N-terminal domain of the glutamate receptor GluA1 [**?**], which exhibits a distinct central channel. The fuzzy-boundary detection method recognizes the channel as a cavity region, where voxels are highly occluded. The boundary of the cavity, and thus its interior, can be established by selecting probability values above a threshold from the 3-D map output by *measure volinterior*.

While computationally more expensive than the standard implementation, the performance of *measure volinterior* with fuzzy-boundary detection is competitive with that of other popular software tools for the analysis of protein cavities. Notably, *measure volinterior* does not require pre-processing of a trajectory to convert it to an anternative format (i.e., PDB). Volumetric analysis for 6500 trajectory frames of the RNA editing ligase 1 (PDB 1XDN) [46], the test system employed for software benchmarks in a recent publication [31], executed on a single thread on a machine containing two Intel Xeon E5-2650 v4 @ 2.20GHz processors using the same parameters from Fig. 5A, requires five minutes of wallclock time with the standard implementation, and 22 minutes with fuzzy-boundary detection enabled. The default behavior for *measure volinterior* is automatic parallelization of ray casting over up to 40 threads, such that the described machine, with 48 total threads, actually runs the benchmark in 3.5 and 9 minutes of wallclock time, respectively. These timings place *measure volinterior* among the fastest reported codes for the examination of cavities [31], even on a desktop workstation. The following section describes deployment of *measure volinterior* on supercomputing architectures to perform large-scale container analyses in a matter of minutes.

### High-Performance Container Analysis

In cases where container trajectories encompass multi-million atom systems over extended simulation timescales, *measure volinterior* is well suited to take advantage of parallel, distributed-memory architectures through VMD’s multiprocessor, GPU-enabled back-end, which can otherwise be bottlenecked by I/O bandwidth in single-node cases. Code Snippet 9 provides a Tcl script demonstrating how to perform container analysis on ORNL’s Titan and Summit and NCSA’s Blue Waters using their high-performance Lustre filesystems and VMD compiled with MPI (see Supplement). With each trajectory frame striped across several object storage targets (OSTs), compute nodes benefit from parallel access, greatly decreasing the burden that I/O places on overall analysis execution time [47]. A schematic illustration of this process is shown in Fig. 6.

**Fig 6.**
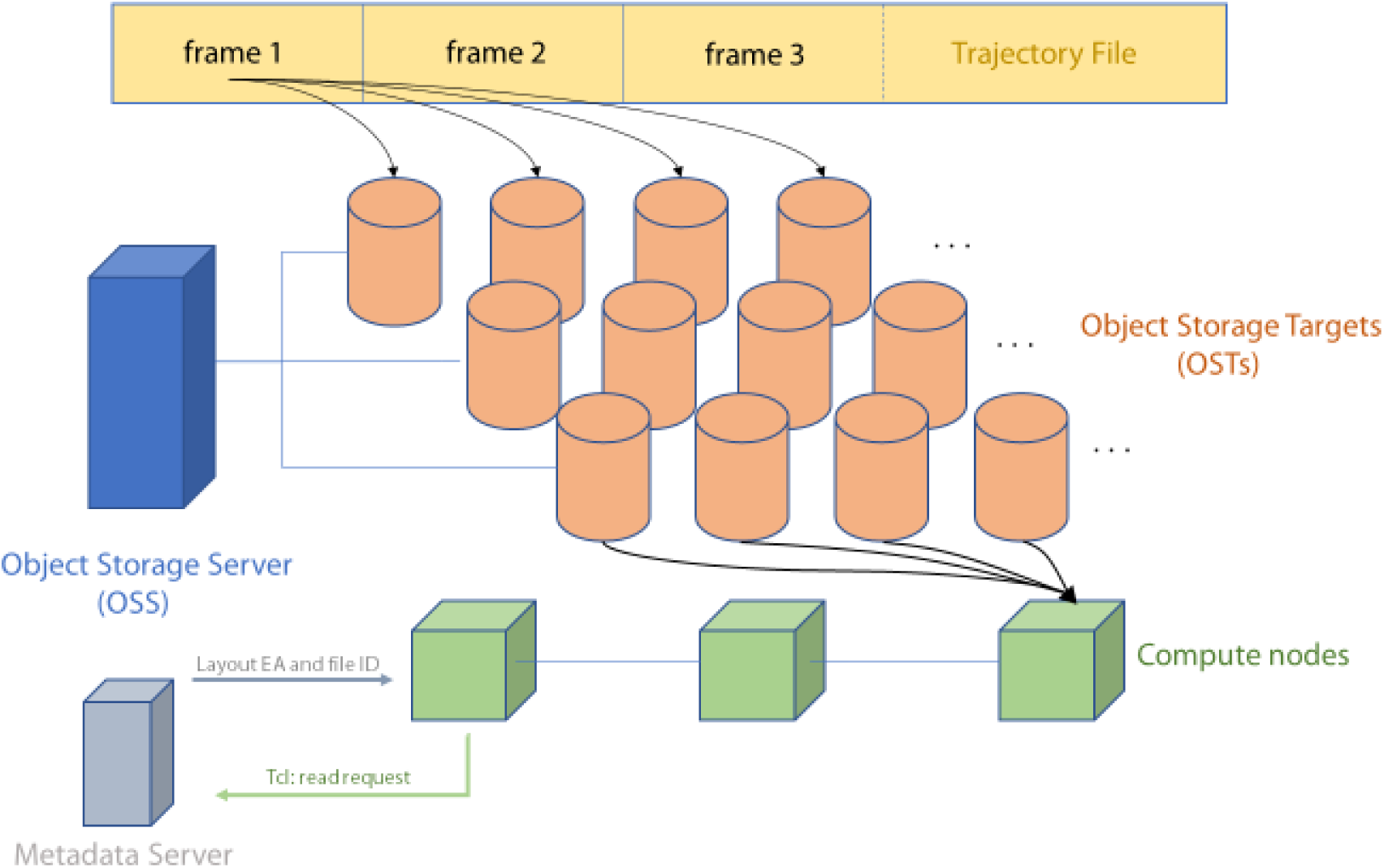
Trajectory frame analysis using the Lustre file system. Each frame in the trajectory file is striped onto separate object storage targets (OSTs). The amount of stripes is dependent on the size of the frame and the available memory in each OST. The object storage server (OSS) interfaces each OST with instructions from the client. Multiple instances of the client are spawned by each compute node once the job is submitted, and each compute node then processes a series of frames using parallel I/O to ameliorate bottlenecking. Instructions are organized through the metadata storage server (MDS), which returns memory addresses and extended attributes in response to high-level read requests sent through Tcl from clients.

As an example, the online storage available to users on NCSA’s Blue Waters supercomputer is comprised of three Lustre storage systems with separate mount points (/scratch, /projects, and /home) that are avaiable from both login and compute nodes. These volumes appear as standard disk partitions in the shell, granting users the capability to interface with the storage as part of a seemingly-standard directory tree, avoiding the operational complexity of the underlying architecture. Once a target directory is created for the input trajectory, it can be striped to a specified count through the command-line.

When running an analysis using a similar framework to the aforementioned, the number of nodes is passed as a user-specified argument to *mpirun* during job submission and is input along with the given trajectory’s framecount to block decompose a set of frames to each specific MPI rank. Each rank runs its own client instance and owns a unique index among all participating nodes. The instance running on node 0 ensures, via a conditional, that rank 0 is responsible for all printing. Once this information is primed for a given node, *parallel barrier* statements instruct each compute node to wait for all of its peers before *measure volinterior* analysis commences.

Using this strategy to perform container analysis on NCSA’s Blue Waters supercomputer, the trajectory for the largest all-atom simulation ever published – an intact HIV-1 capsid in explicit solvent, encompassing 64 million atoms simulated over 1.2 microseconds and boasting a data storage footprint of 70 TB [4] – can be processed in under ten minutes of wallclock time. HIV-1 capsid trajectory analysis employed a stripe count of 120 and a parallel load of 12 frames per compute node.

## Conclusion

The *measure volinterior* command implemented in VMD 1.9.4 represents a marriage between VMD’s established molecular visualization and high-performance analysis features to realize a brand new capability: 3-D ray casting allows rapid classification of the space surrounding biomolecular containers regardless of container shape and constituent particle count, thereby enabling examination of containers on a scale that was previously unfeasible. Biomolecular containers are essential biological machines, whose study is critical to understanding the living cell. Containers such as enveloped virions [2, 3], organelles [1, 48], and potentially even minimal cells will no doubt be among the major systems investigated with MD simulations on upcoming exascale supercomputing platforms, which will enable calculations encompassing billions of particles. Beyond enabling new scientific discoveries, large-scale MD simulations will continue to drive the development of novel technologies to overcome the challenges of working with big datasets [4, 49]. High-performance analysis software that can efficiently characterize the properties of biomolecular containers, particularly those that support tracking of solvent molecules over many trajectory frames, will drive the revelation of important discoveries arising from container simulations. With the new *measure volinterior* capability, VMD will enable researchers to routinely employ sophisticated biomolecular container analysis in the next supercomputing era.

## Acknowledgments

We acknowledge funding by the National Institutes of Health grants P50GM082251, P30GM110758-04, and P41GM104601 and the University of Delaware. This research is part of the Blue Waters sustained-petascale computing project, which is supported by the National Science Foundation (awards OCI-0725070 and ACI-1238993) and the state of Illinois. This work also used the Extreme Science and Engineering Discovery Environment (XSEDE), which is supported by National Science Foundation grant number ACI-1548562. An award of computer time was also provided by the Innovative and Novel Computational Impact on Theory and Experiment program (INCITE award BIO024). This research also used resources from the Oak Ridge Leadership Computing Facility (OLCF) at Oak Ridge National Laboratory, which is supported by the Office of Science of the Department of Energy under Contract DE-AC05-00OR22725. We acknowledge a Director’s Discretionary award on the Summit supercomputer from the OLCF. Early VMD software development access to the Summit supercomputer was provided as part of the Center for Accelerated Application Readiness project at the OLCF.

## Author Contributions

J.R.P and J.S. designed the research. A.B., J.H., J.S. and J.R.P. performed molecular modeling. A.B., J.H., J.S. and J.R.P wrote *measure volinterior* code. A.B., J.H., J.S. and J.R.P. validated results from the accelerated code. A.B., J.H., J.S. and J.R.P. wrote the manuscript.

## Supporting Information

**Fig S1.**
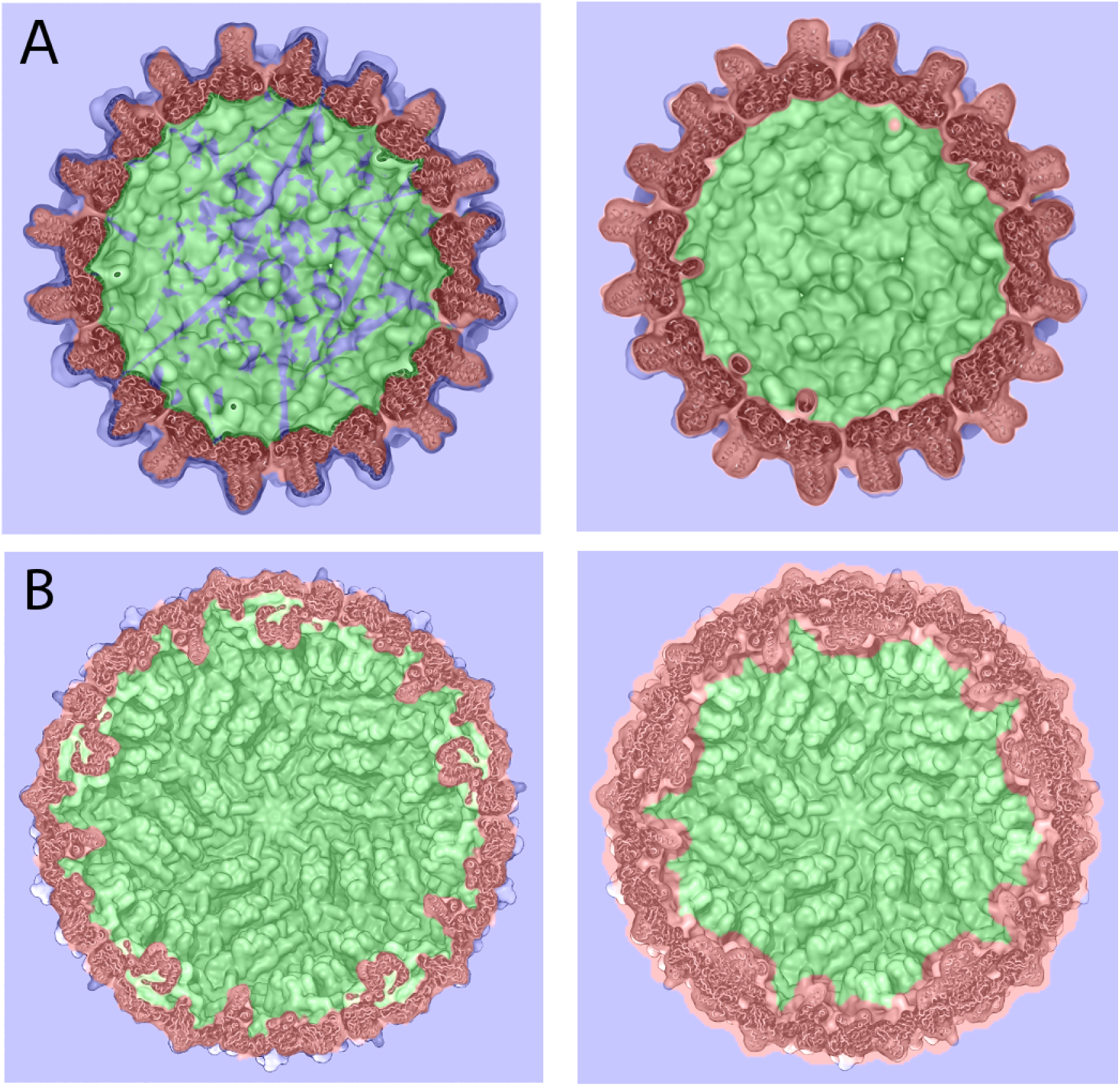
Effect of *measure volinterior* parameters on quality of molecular surfaces and space classification. (A) Using parameters that fail to mask container openings over all frames of a simulation trajectory leads to inaccurate classification of the container’s interior space. Although the *measure volinterior* parameters used to analyze the experimental HBV capsid structure produced a continous molecular surface without holes (Fig. 2B), the same parameters applied to analyze a frame extracted from an MD simulation of the capsid at 310 K [5] allow the escape of rays due to fluctuations of capsid pores (left). Using parameters that produce a slightly bulkier molecular surface successfully masks the pore openings over all trajectory frames and ensures more accurate space classification (right). (B) Using parameters that produce bulky molecular surfaces reduces the level of topological detail captured for the container and leads to less accurate classification of its surrounding space. The *measure volinterior* parameters used to analyze the experimental structure for the ZIKV E/M protein shell yield a space classification map that closely tracks the container’s surface features (left). In contrast, parameters that produce a more bulky molecular surface yield a map that assigns additional space around the proteins as part of the container itself, leading to a less accurate representation of the container’s internal volume (right)

**Fig S2.**
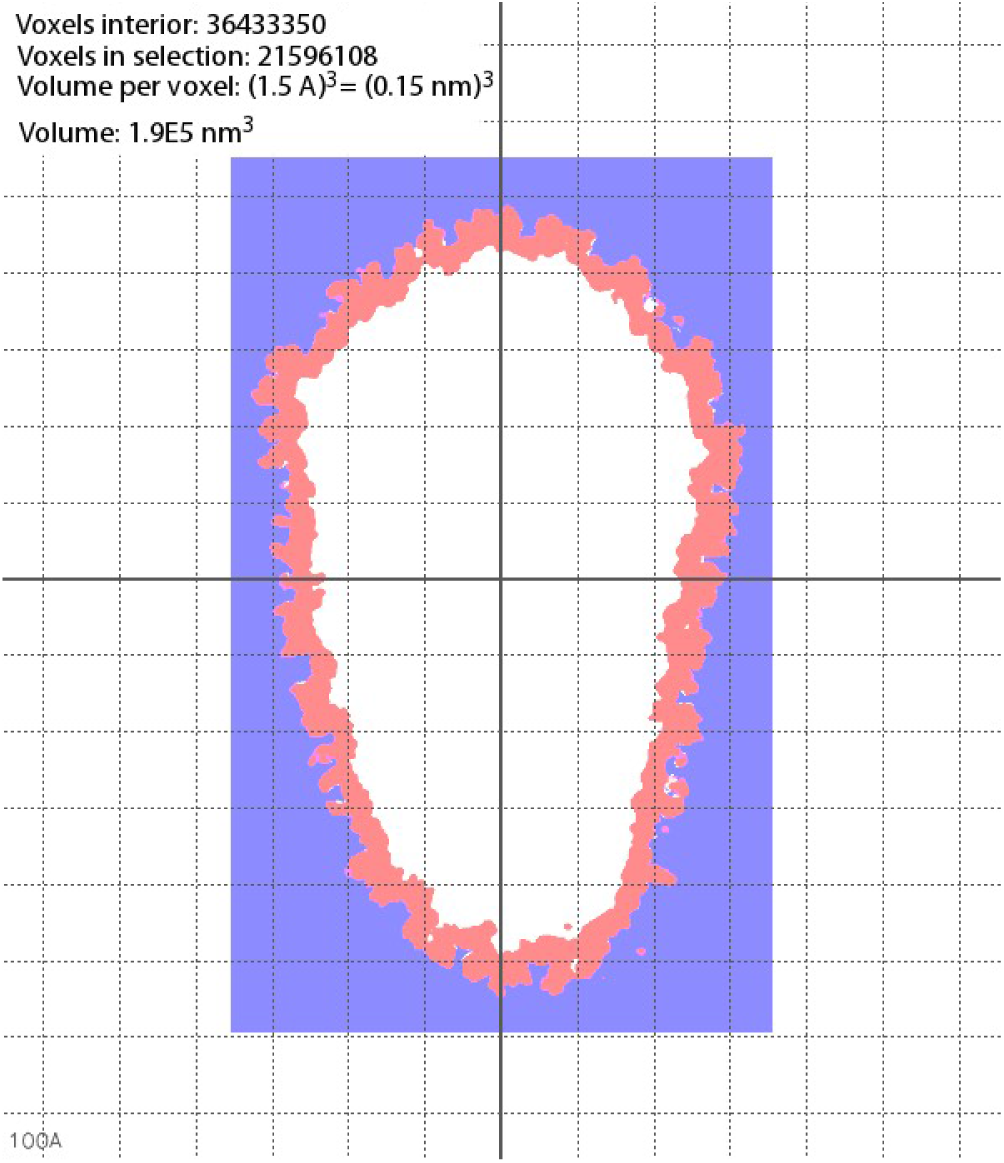
Volume of the HIV-1 capsid. Using *measure volinterior*, the volume of the interior of an irregular container can be accurately calculated. For PDB 3J3Q [15] using the parameters given in Table 1, the internal volume is 1.9 × 10^5^ nm^3^.

**Code Snippet 7.**
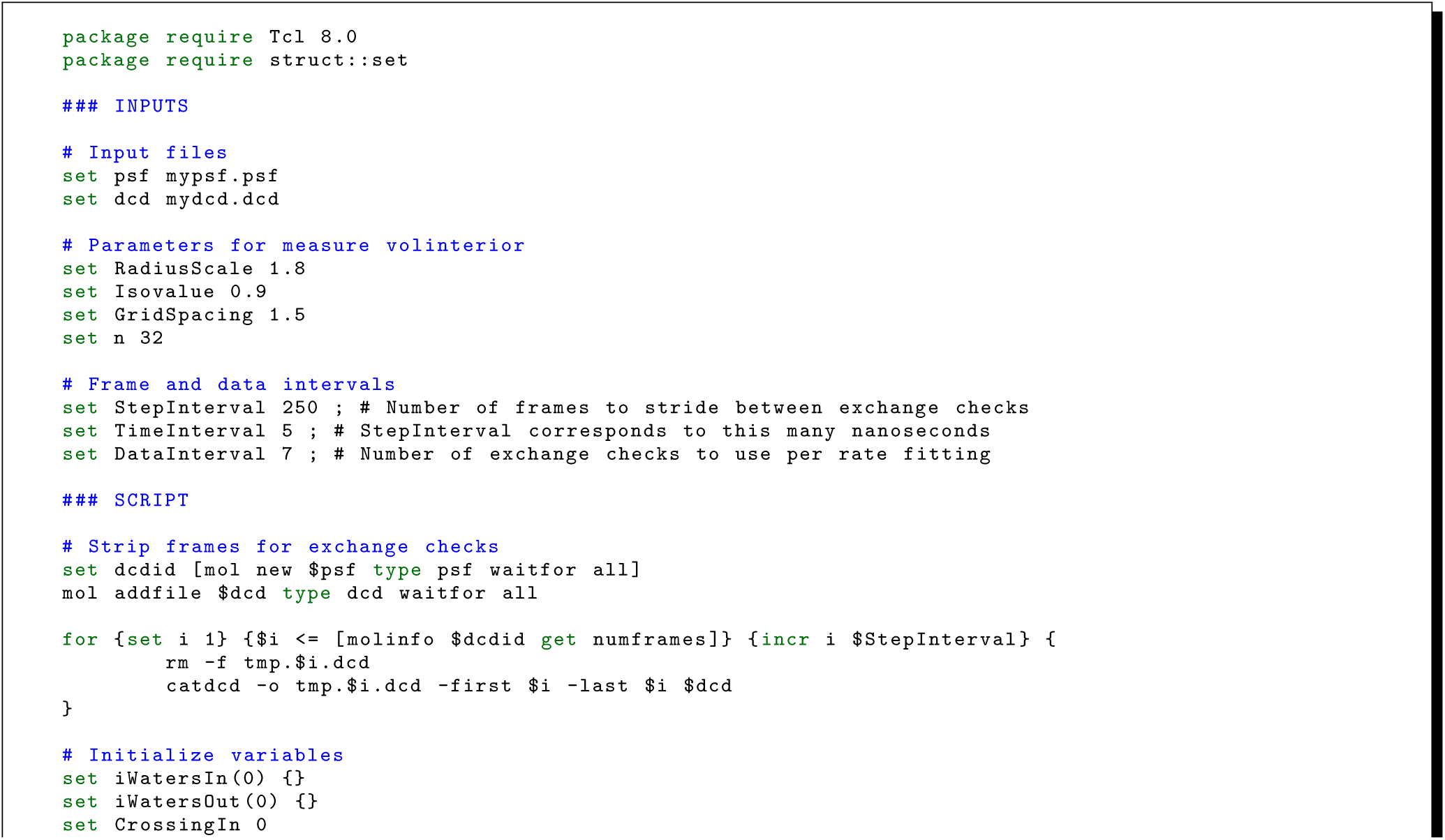
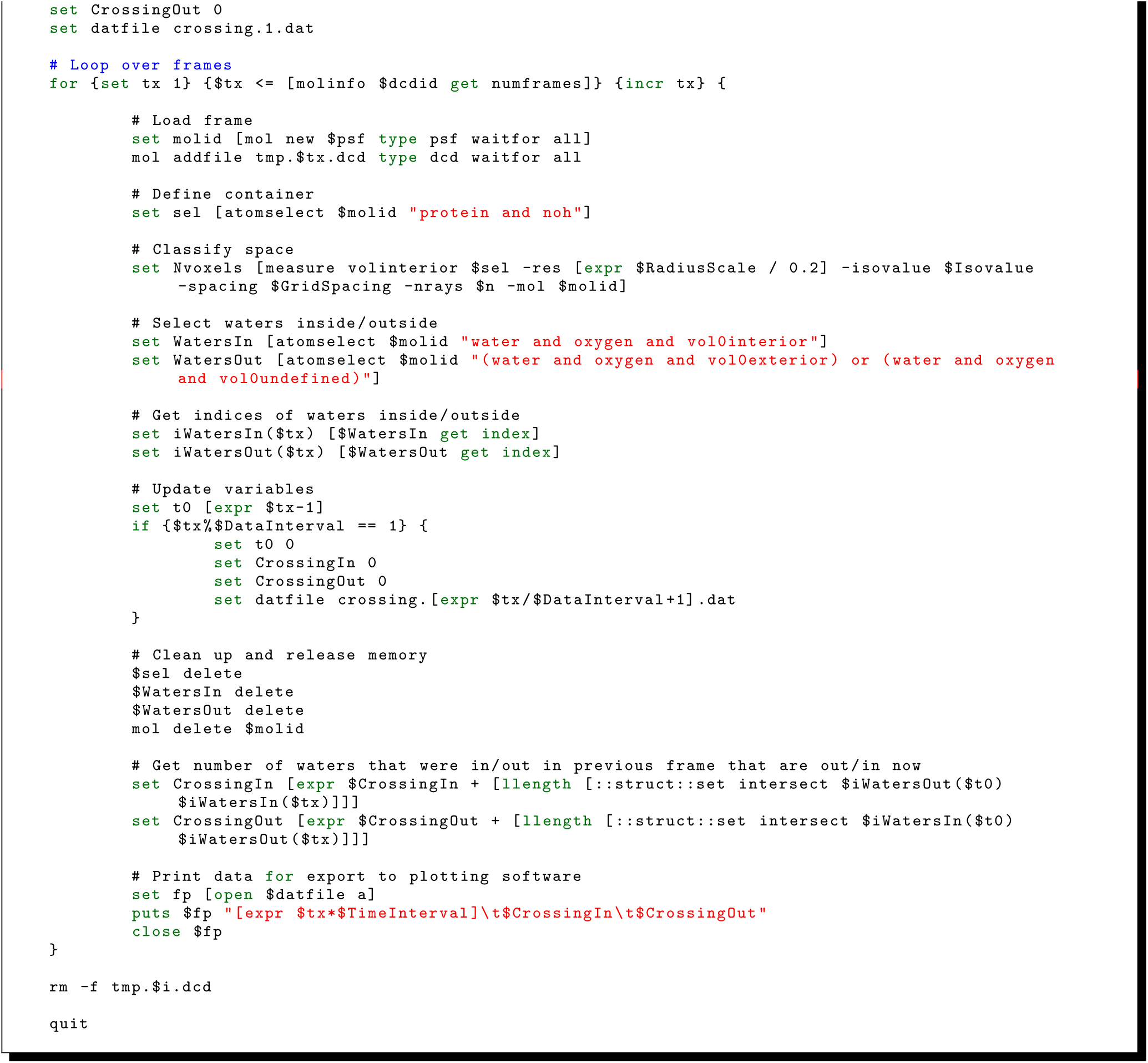
Extracting data to calculate water exchange rates using *measure volinterior*.

**Code Snippet 8.**
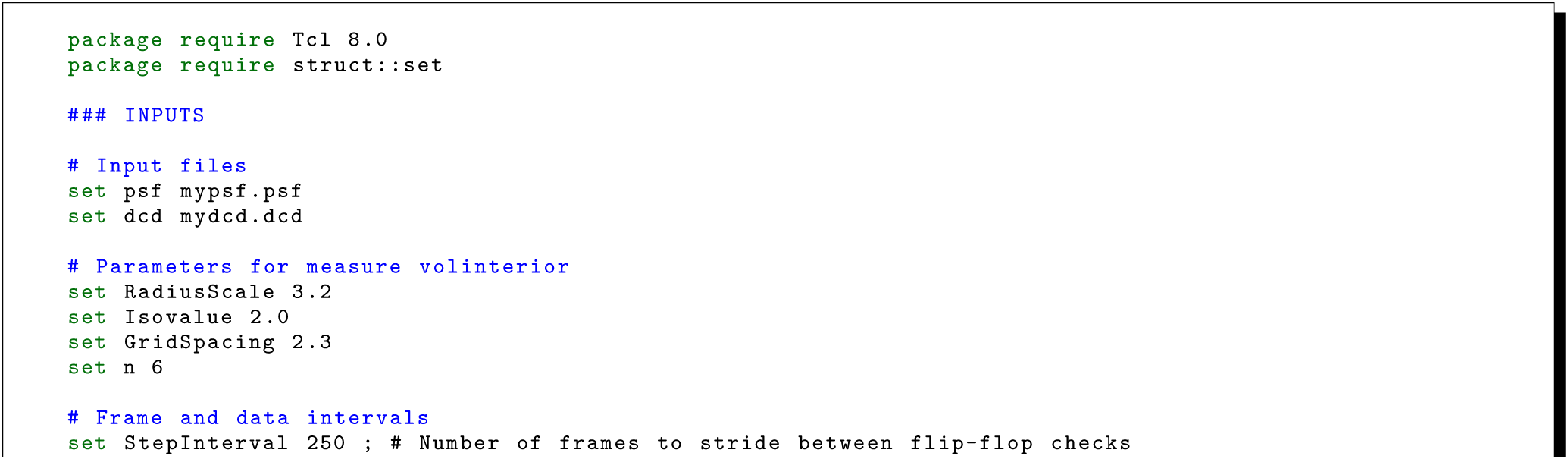
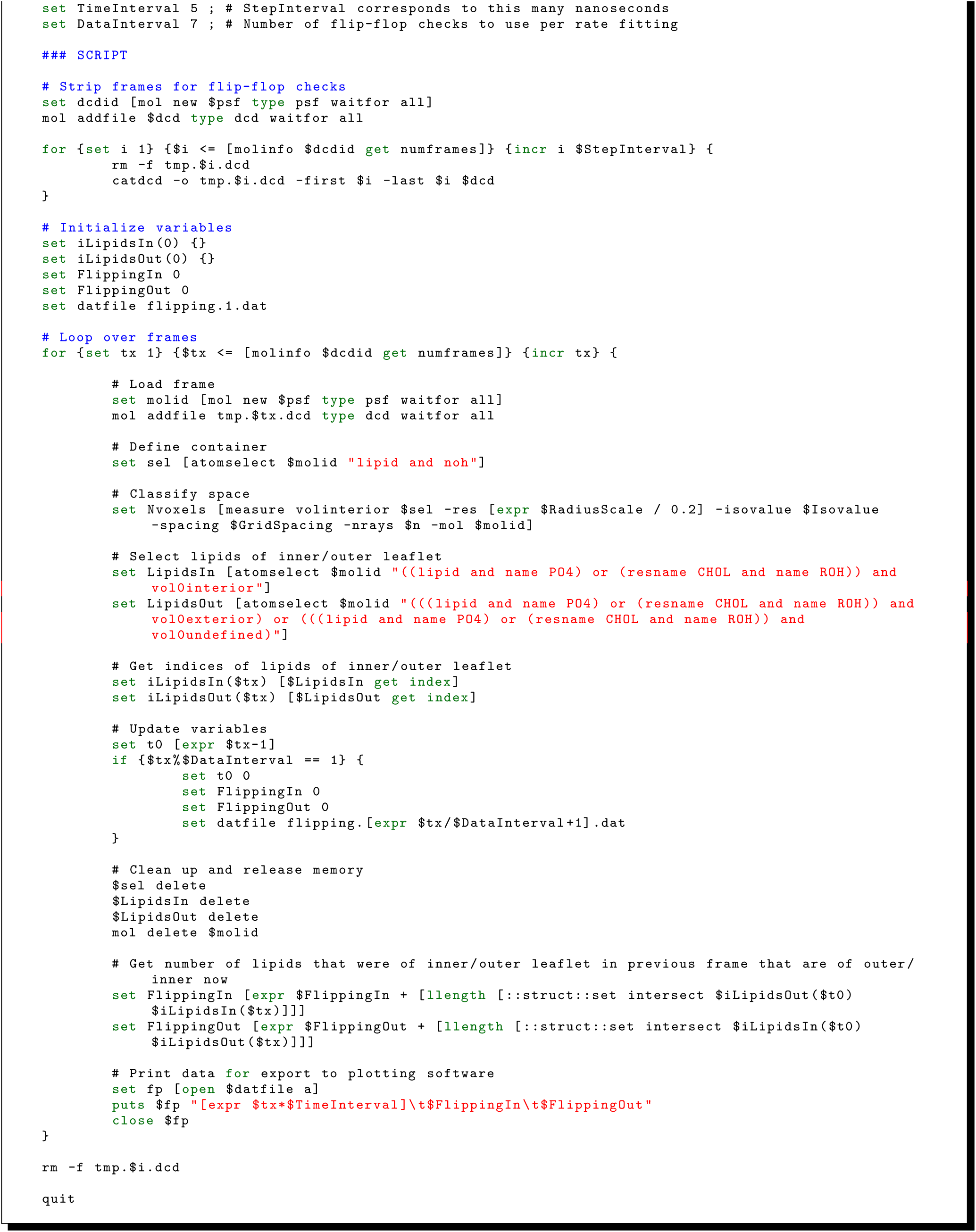
Extracting data to calculate lipid transverse diffusion rates using *measure volinterior*.

**Code Snippet 9.**
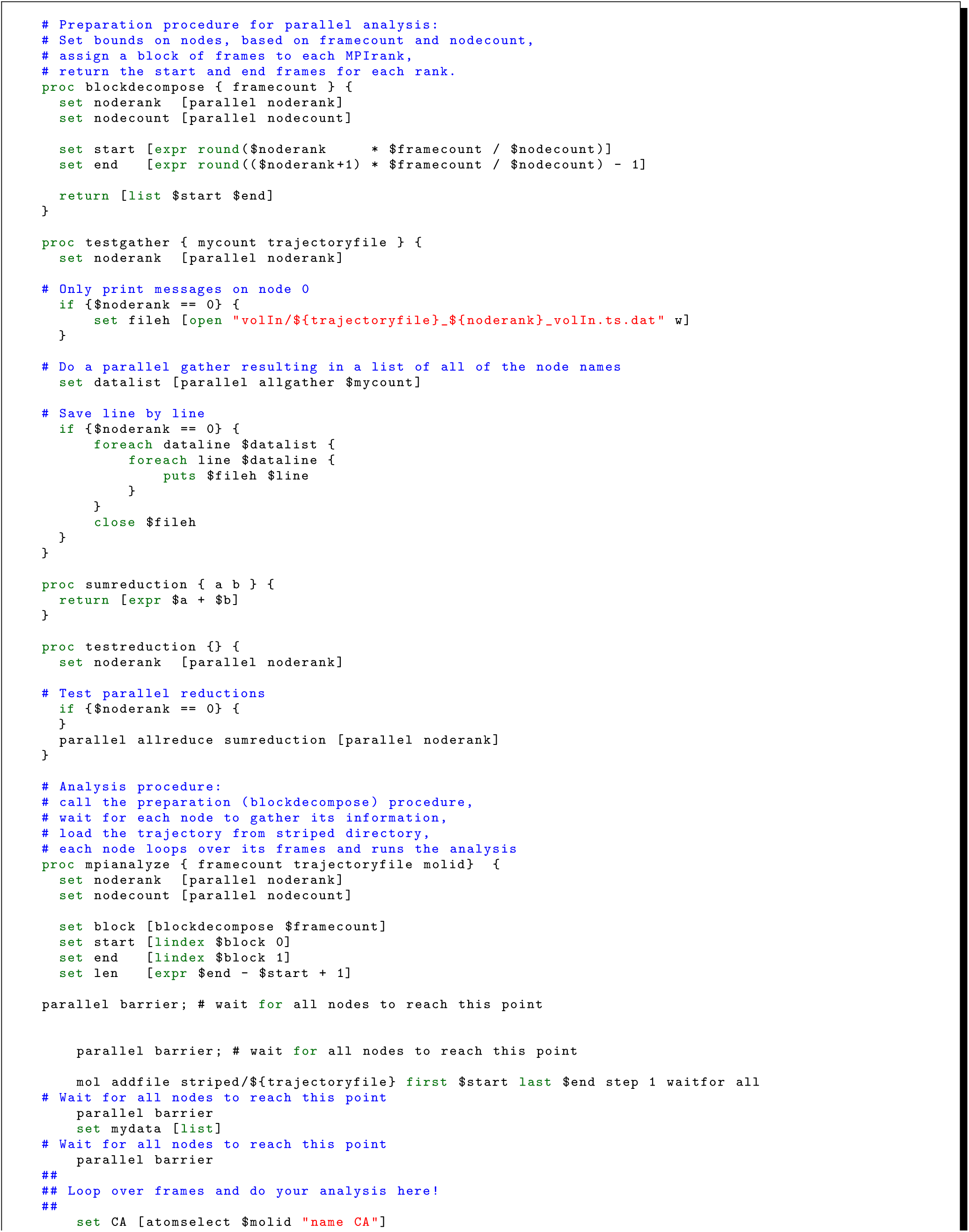
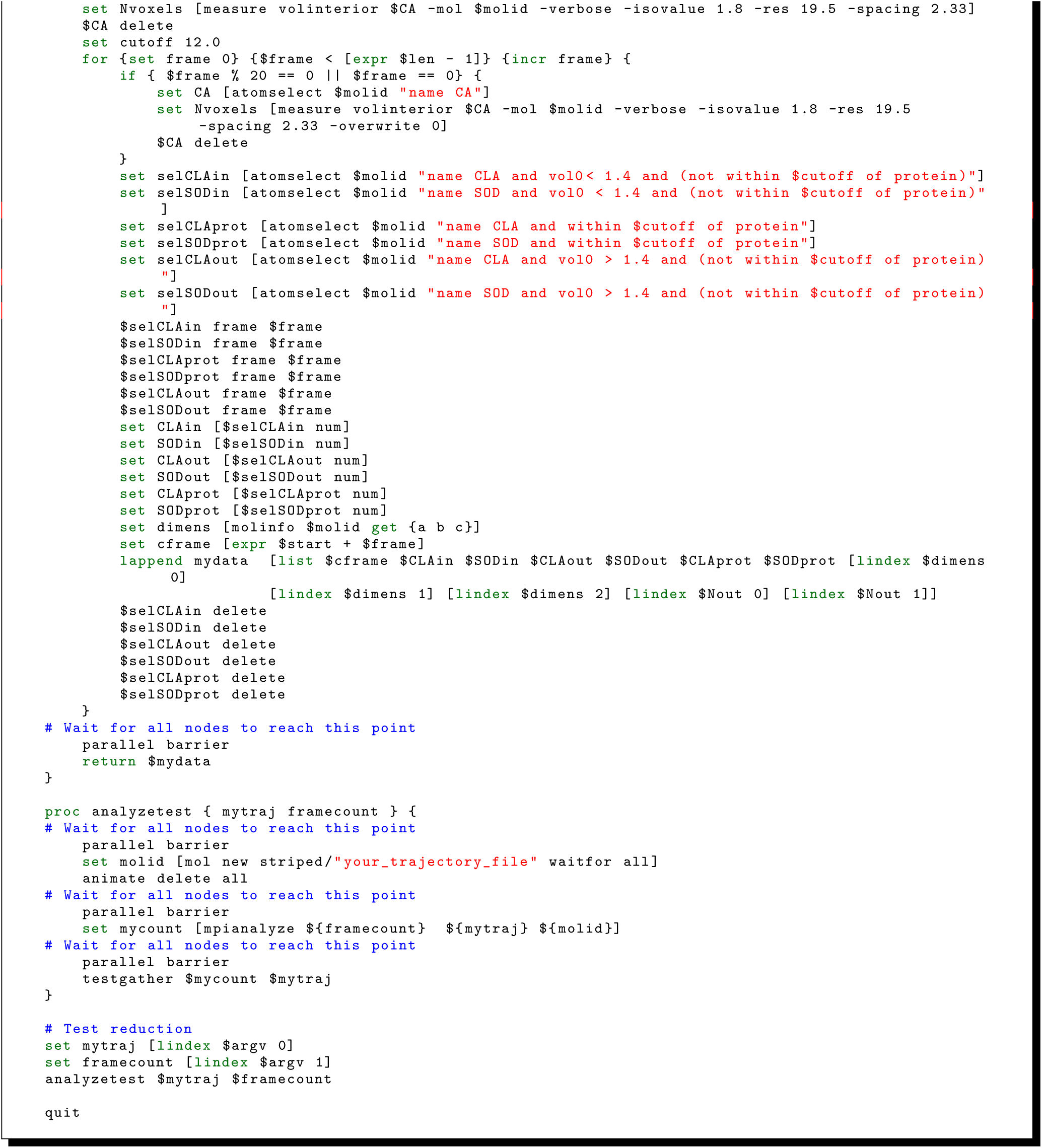
High-performance analysis of particle densities using VMD compiled with MPI.

